# SETMAR functions in illegitimate DNA recombination and non-homologous end joining

**DOI:** 10.1101/465138

**Authors:** Michael Tellier, Ronald Chalmers

## Abstract

In anthropoid primates, SETMAR is a fusion between a methyltransferase gene and a domesticated DNA transposase. SETMAR has been found to be involved in several cellular functions including regulation of gene expression, DNA integration and DNA repair. These functions are thought to be mediated through the histone methyltransferase, the DNA binding and the nuclease activities of SETMAR. To better understand the cellular roles of SETMAR, we generated several U2OS cell lines expressing either wild type SETMAR or a truncated or mutated variant. We tested these cell lines with *in vivo* plasmid-based assays to determine the relevance of the different domains and activities of SETMAR in DNA integration and repair. We found that expressing the SET and MAR domains, but not wild type SETMAR, partially affect DNA integration and repair. The methyltransferase activity of SETMAR is also needed for an efficient DNA repair whereas we did not observe any requirement for the putative nuclease activity of SETMAR. Overall, our data support a non-essential function for SETMAR in DNA integration and repair.

## Introduction

SETMAR is an anthropoid primate-specific fusion between a histone methyltransferase gene, connected to dimethylation of histone H3 lysine 36 (H3K36me2), and a domesticated Hsmar1 transposase (1-3). The transposase domain is 94% identical to the Hsmar1 transposase consensus sequence but three mutations, including the DDD to DDN mutation of the catalytic triad, completely abolish the transposition activity of SETMAR (4-6). Although some activity, particularly 5’-end nicking, is recovered *in vitro* in the presence of DMSO and Mn^2+^, it is likely to not be significant in physiological conditions (4). Nevertheless, the transposase DNA-binding domain of SETMAR is under purifying selection and retains robust transposon end binding and the ability to form dimer (2, 4, 7). It has recently been shown that SETMAR could regulate gene expression in human cells through the combination of its binding to the Hsmar1 transposon ends scattered in the human genome and its methyltransferase activity (7).

Earlier experiments with SETMAR revealed that it was involved in illegitimate DNA integration and DNA repair through the non-homologous end joining repair (NHEJ) pathway (1, 8). NHEJ is one of the four pathways used by the cell to repair DNA double-strand breaks (DSBs) and the primary repair pathway throughout the cell cycle (9). NHEJ is a template-independent DNA repair pathway, which relies on Ku proteins to bind the DNA free ends, on nucleases, such as Artemis, or polymerases to trim or fill the DNA overhangs and on the DNA ligase IV complex to ligate together the two blunt ends (9).

Illegitimate DNA recombination and lentivirus cDNA integration are dependent of the NHEJ pathway but the mechanism responsible for plasmid integration, which cannot rely on an integrase, remains uncertain (10, 11). The current model states that the circular plasmid needs to be linearized by a DSB for recruiting DNA repair proteins on the plasmid ends. For genomic integration to happen, one plasmid end needs to be in the vicinity of a genomic lesion for the NHEJ proteins to use the linearized plasmid DNA to repair the genomic DSB (11).

One of the difficulties in understanding the functions of SETMAR in DNA repair is that it produced a response in a number of different assays, suggesting that it was involved in many different aspects of DNA metabolism. For example, its overexpression promotes classical NHEJ, the random integration of transfected plasmid DNA and the restart of stalled replication forks (1, 12). Based on *in vitro* analysis, it has been hypothesized that purified SETMAR could act as an endonuclease like Artemis (13, 14). However, SETMAR endonuclease activity has only been established *in vitro* and recent papers question its relevance *in vivo* (14, 15). In contrast to Artemis, which promotes both trimming of DNA overhangs and DNA repair in cell extract assays, SETMAR did not stimulate DNA repair and only promotes trimming in one assay.

The SET methylase-domain of SETMAR was shown to interact with PRPF19, also known as PSO4, which is a protein involved in the classical NHEJ and the spliceosome, and with DNA ligase IV, which is responsible for ligating the blunt ends in NHEJ (16, 17). The interaction with PRPF19 was predicted to target SETMAR to double strand DNA breaks where the SET domain could dimethylate the histone H3 lysine 36 of neighbouring nucleosomes (18). This epigenetic mark recruits and stabilizes the anchoring of Ku70 and NBS1, two early acting NHEJ factors, to the DNA ends (18). Two other papers linked the increase in H3K36me2 following DSBs to the inhibition of KDM2A and KDM2B, two histone demethylases involved in the removal of H3K36 methylation (19, 20). However, a recent study did not observe an increase in H3K36me2 around DSB sites (21).

To better understand the functions of SETMAR in NHEJ, we produced several U2OS cell lines expressing different SETMAR constructs to test the role of the SET and MAR domains and the methyltransferase, DNA binding and nuclease activities of SETMAR in illegitimate DNA integration and repair. We found that expression of the SET and MAR domains, but not of wild type SETMAR, affect DNA integration and repair. SETMAR methyltransferase activity is required for an efficient DNA repair but we did not observe any role for the putative nuclease activity of SETMAR. We hypothesize that the dimerization of SETMAR imposed by the MAR transposase domain could have interfere with the pre-fusion functions of the SET domain.

## Materials and Methods

### Media and growth conditions

The T-Rex-U2OS cell lines were maintained in complete Dulbecco’s modified Eagle’s medium (DMEM, Sigma) supplemented with 10% heat inactivated Foetal Bovine Serum (FBS), 100 u/ml of streptomycin, 100 μg/ml of penicillin, and 5 μg/ml of blasticidin at 37°C with 5% CO_2_. The medium of T-Rex-U2OS cell lines stably expressing a gene of interest from an integrated pcDNA4TO plasmid was supplemented with 400 μg/ml of zeocin.

### Plasmids

An artificial codon-optimized version of SETMAR was synthesized by Gene Art (Thermo Fischer) and cloned into pcDNA4TO at the EcoRI/NotI restriction sites. The truncated and mutant (N210A, R432A and D483A) versions of SETMAR were produced by PCR. pRC1712 was constructed by cloning a neomycin resistance gene into pBluescript SKII+ (Agilent) at the BamHI restriction site.

### Stable transfection of T-Rex-U2OS cells

For each transfection, 2.5×10^5^ of cells were seeded in a 6-well plate and grown overnight in DMEM supplemented with 10% FBS. The plasmids were transfected using Lipofectamine 2000 (Invitrogen), following manufacturer’s instruction. After 24 hours, a quarter of the cells were transferred to 100 mm dishes and the medium supplemented with 400 μg/ml of zeocin (Invivogen). After 2 weeks of selection, single foci were picked and grown in a 24-well plate. The expression of the gene of interest was verified in each cell line by inducing the PCMV promoter with doxycycline at a final concentration of 1 μg/ml for 24 hours. The list of cell lines used in this study is presented in Table 1.

**Table 1:**
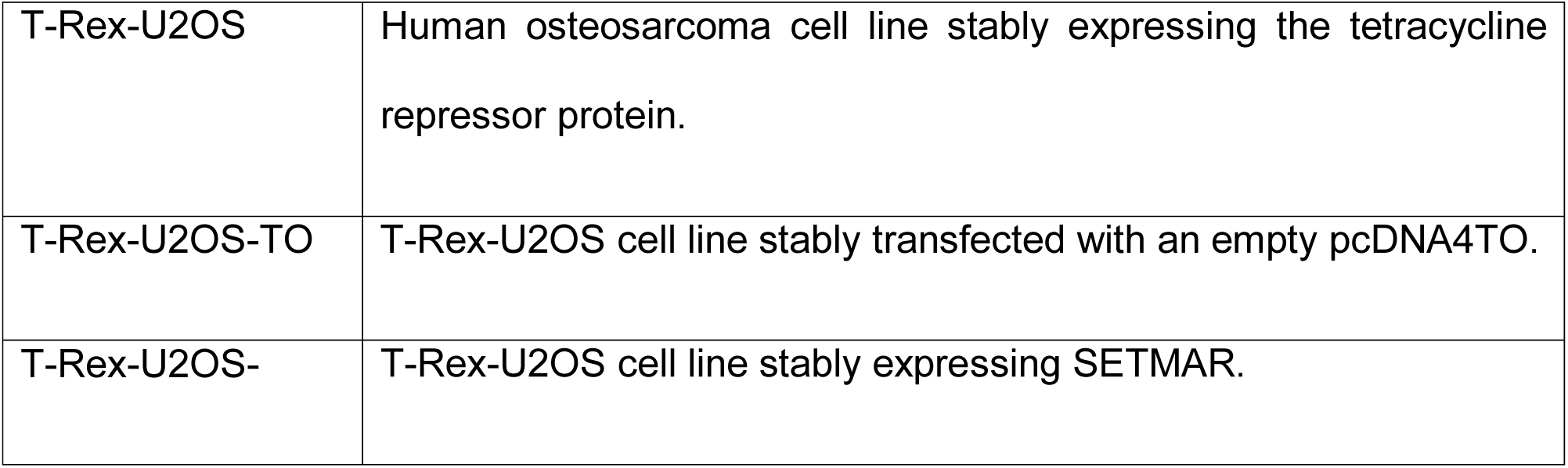
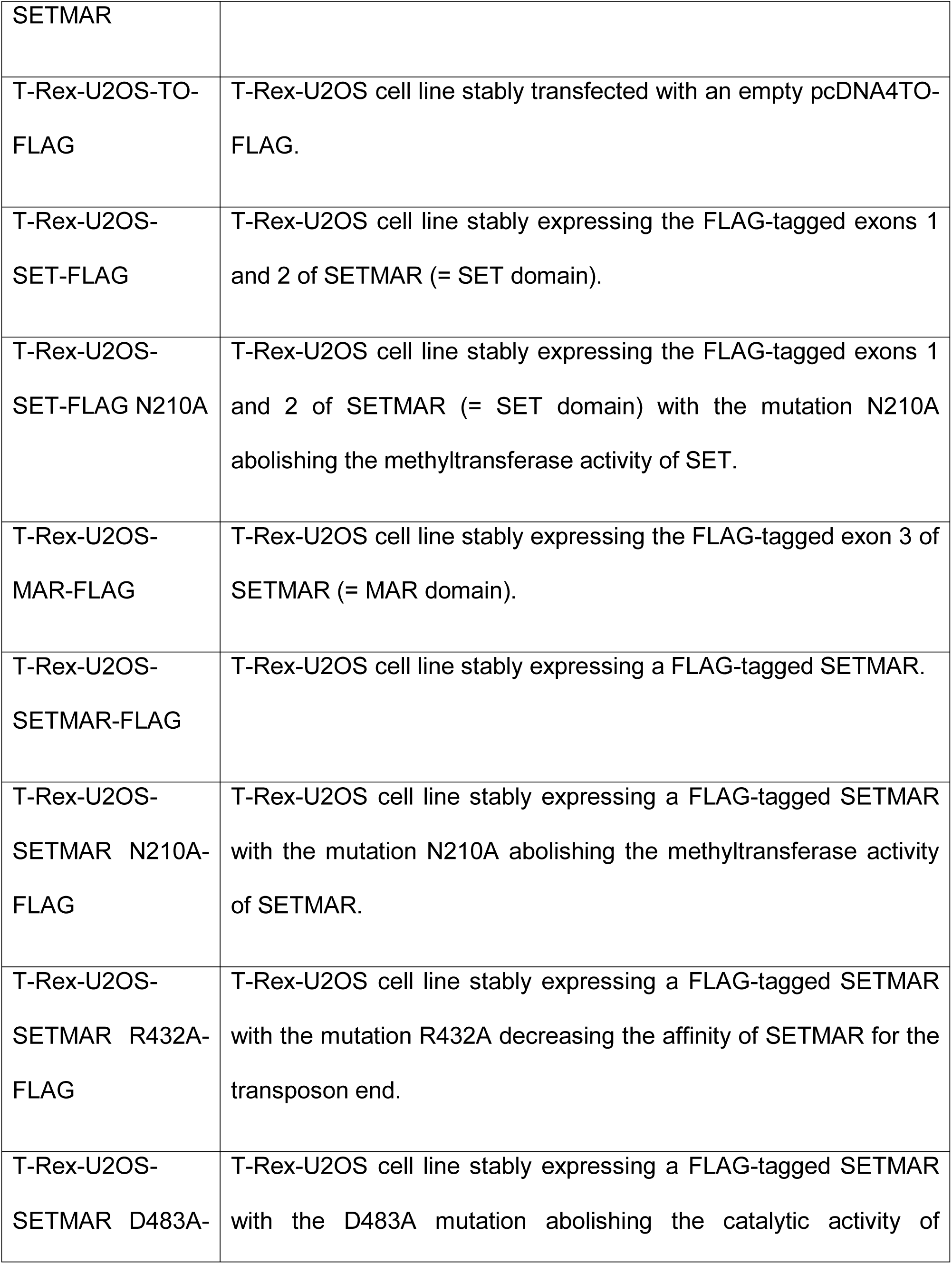
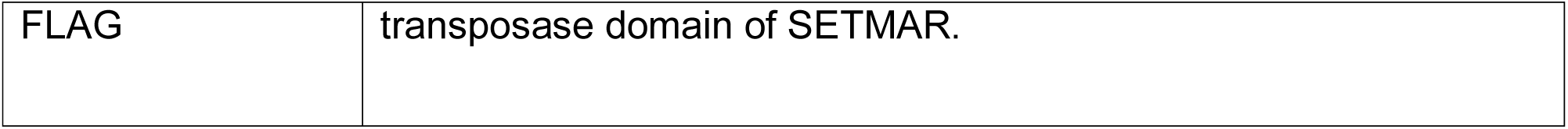
Mammalian cell lines used in this study

### Western blotting

Whole cell extracts were harvested from cultures at ~90% confluency in six-well plates. Briefly, cells were washed two times with ice-cold PBS then pelleted for 5 minutes at 3000 x g at 4°C. Samples were resuspended in 100 μl of Radio ImmunoPrecipitation Assay (RIPA) buffer (10 mM Tris-HCl pH8.0, 150 mM NaCl, 1 mM EDTA, 0.1% SDS, 1% Triton X-100, 0.1% sodium deoxycholate) with freshly added protease inhibitor cocktail (Roche Applied Science) and incubated on ice for 30 minutes, with a vortexing every 10 minutes. Cell lysates were centrifuged for 15 minutes at 14000 x g at 4°C and the protein in the supernatants was quantified by the Bradford assay.

For each western blot, 20 μg of proteins were mixed with 2X SDS loading buffer, boiled for 5 minutes, and electrophoresed on a 10% SDS-PAGE gel. Proteins were transferred to a polyvinylidene difluoride (PVDF) membrane, which was blocked in 5% milk or BSA (Roche) and incubated with specific primary antibodies at 4°C overnight. After washing, membranes were incubated with horseradish peroxidase (HRP)-conjugated secondary antibodies for one hour at room temperature, washed, and signals were detected with the ECL system (Promega) and Fuji medical X-ray film (Fujifilm).

The following antibodies were used: anti-beta Tubulin (rabbit polyclonal IgG, 1:500 dilution, ab6046, Abcam), anti-Hsmar1 antibody (goat polyclonal, 1:500 dilution, ab3823, Abcam), anti-FLAG (rabbit, 1:500 dilution, F7425, Sigma). The secondary antibodies were horseradish peroxidase-conjugated anti-goat (rabbit polyclonal, 1:5000 dilution, ab6741, Abcam) and anti-rabbit (goat polyclonal, 1:5000-1:10000, ab6721, Abcam).

### Growth rate

At day 0, 2×10^4^ cells were seeded in eight 6 cm dishes for each cell line and one dish was count every day for eight days using a hemocytometer.

### Illegitimate DNA integration assay

For integration assays in the T-Rex-U2OS cell lines, 8×10^5^ cells were seeded onto 6-well plates with 2.5 μg of circular or linearized pRC1712 and 5 μl of Lipofectamine 2000 (Invitrogen). Twenty-four hours later, cells were trypsinized and 5×10^4^ cells of each transfection were seeded onto 10 cm dishes in medium containing 800 μg/ml of G418 (Sigma). After two weeks of selection, surviving foci were fixed for 15 min with 10% formaldehyde in PBS, stained for 30 min with methylene blue buffer (1% methylene blue, 70% ethanol), washed with water, air dried, and photographed. The transfection efficiency was tested by transfecting a pEGFP plasmid. After 24 hours, the live cells were observed using a Carl Zeiss Axiovert S100 TV Inverted Microscope with an HBO 100 illuminator. The transfection efficiency was found to be similar between the different cell lines.

### Non-homologous end-joining assay and FACS analyses

Prior to transfection, the pEGFP-Pem1-Ad2 plasmid was digested overnight with *HindIII* or *I-SceI*. The digested plasmids were heat-inactivated and column-purified before being co-transfected with a pRFP plasmid for controlling the transfection efficiency. A day before transfection, 8×10^5^ cells were seeded in 60 mm dishes for obtaining a ~70 % confluency on the transfection day. Transfections were performed with 3 μg of linear pEGFP-Pem1-Ad2, 3 μg of pRFP and 14 μl of Lipofectamine 2000 (Invitrogen), according to manufacturer’s instructions. After 24 hours, green (GFP) and red (RFP) fluorescence was measured by fluorescence-activated flow cytometry (FACS). For FACS analysis cells were harvested with Accutase (Sigma), washed once in 1X PBS and fixed in 2% formaldehyde (Sigma). FACS analysis was performed on a Coulter FC500 (Beckman Coulter). The numbers of repaired events are reported as the ratio of green and red positive cells over the total number of red positive cells. This ratio normalizes the numbers of repaired events to the transfection efficiency. The values for all the cell lines are reported as a percent of the control cell lines.

## Results

### SETMAR overexpression does not promote cell proliferation in the U2OS cell line

It was previously observed that SETMAR overexpression increases the growth rate of the HEK293 and HEK293T cell lines (22). Conversely, SETMAR depletion by RNA interference or CRISPR/Cas9 knock-out was found to decrease the growth rate of THP-1 and DLD-1 cancer cells, respectively (23, 24). We previously shown that a U2OS cell line mildly overexpressing SETMAR that SETMAR is involved in the regulation of the expression of a broad set of genes (7). However, we did not found an enrichment for genes involved in cell cycle (7). To determine whether altering SETMAR expression level also affects the growth rate of the U2OS cell line, we tested three stable T-Rex-U2OS cell lines overexpressing at different level Flag-tagged version of SETMAR and one cell line expressing the SET domain only (Fig 1A). The expression level of the SET domain or SETMAR was determined by western blotting using an anti-FLAG antibody to allow the comparison between the cell lines. The growth rate was determined by counting the number of cells across a period of eight days (Fig 1B). A small but significant decrease in cell proliferation was observed for most of the cell lines overexpressing SET or SETMAR after 5 to 6 days.

**Fig 1.**
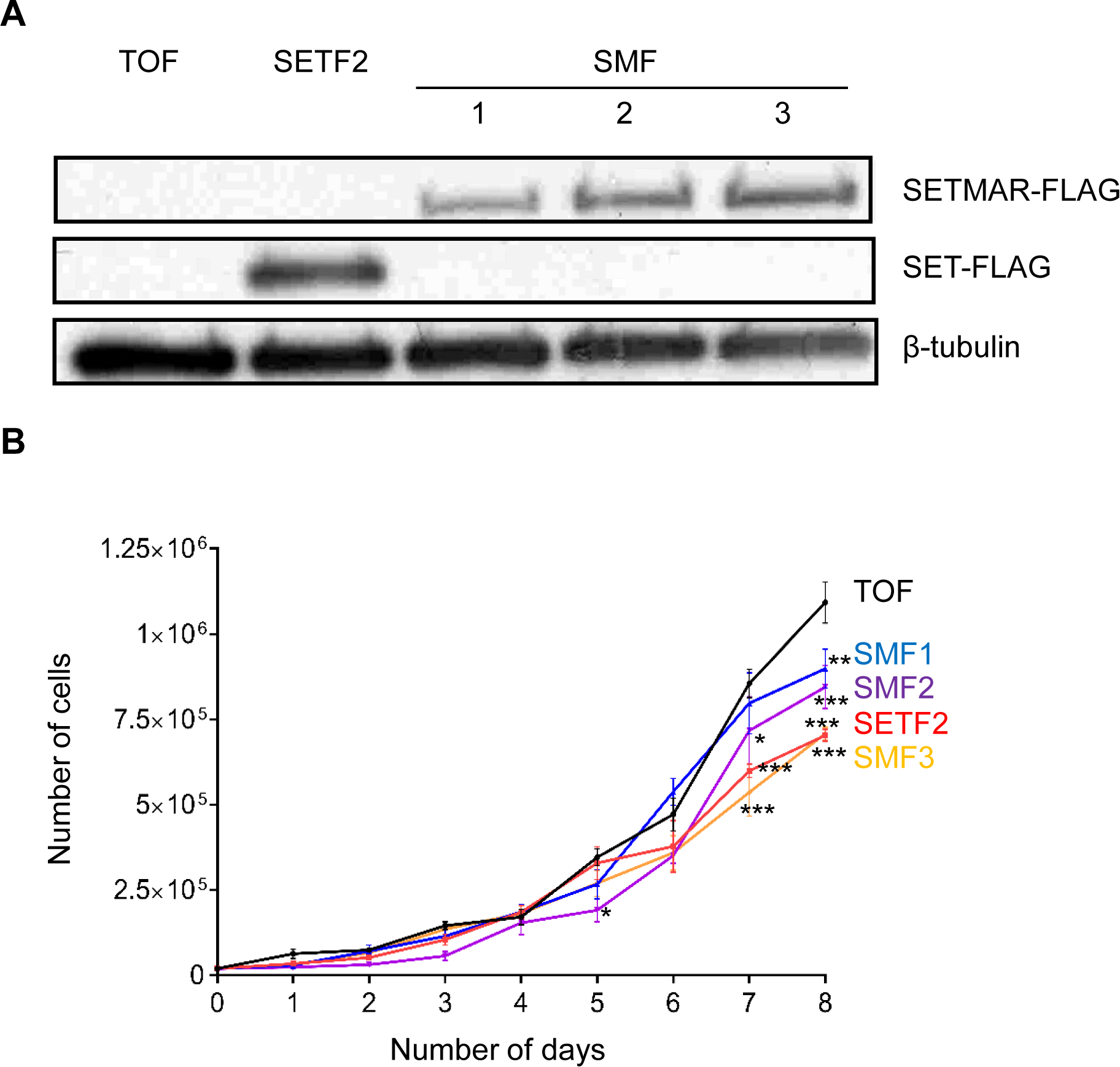
The overexpression of SET or SETMAR do not promote cell proliferation in an U2OS genetic background. **A**, Western blot for the FLAG-tagged SETMAR in the U2OS, SETF and SMF cell lines. The western blot was performed with anti-FLAG and anti-β-tubulin antibodies. **B**, Growth rate of U2OS, SETF and SMF cell lines. At day 0, 2.0×10^4^ cells were seeded in eight dishes and one dish was counted every day for eight days. Average ± S.E.M. of 3 to 5 biological replicates. Statistical test: t-test with Holm-Sidak correction, ^∗^ p-value < 0.05, ^∗∗^ p-value < 0.01, ^∗∗∗^ p –value < 0.001

### Characterization of the different SETMAR constructs

To improve our understanding of SETMAR roles in illegitimate DNA integration and the NHEJ pathway, we produced several U2OS cell lines stably overexpressing wild type, truncated or mutant version of SETMAR (Fig 2A and Table 2). SETMAR probably functions as a dimer in the cell with the transposase domain providing the whole dimer interface (25). The endogenous concentration of SETMAR in the U2OS cell line is low, with less than 500 molecules per cell (26). The overexpression of a SETMAR mutant should therefore produce dimers of two mutant monomers and dimers containing wild type and mutant monomers. Dimers with two wild type monomers are the less likely so SETMAR activity in the cell should be hindered by the overexpression of the mutants. The expression level of each cell line was determined by western blotting using anti-SETMAR and anti-FLAG antibodies (Fig 2B). An anti-FLAG antibody was used for the cell lines containing an F (for FLAG-tag) in their names. SM2 and 3 overexpress a version of SETMAR without any FLAG-tag so an antibody against the last nine amino acids of SETMAR was used to determine their expression level.

**Fig 2.**
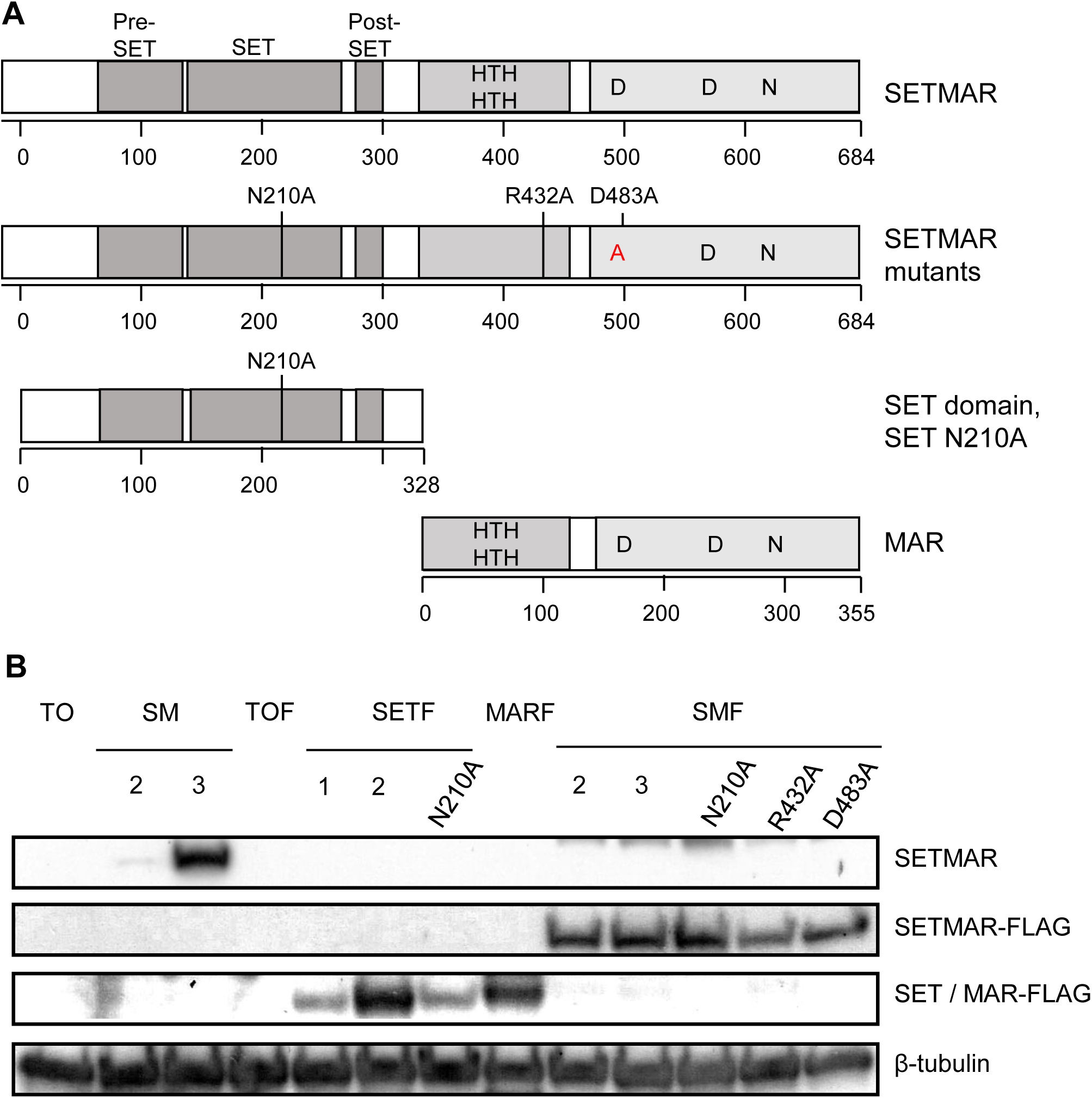
U2OS cell lines used in the *in vivo* DNA repair assay. **A**, Schematic representation of SETMAR, SET and MAR and the location of the different mutations. **B**, Western blot for the FLAG-tagged SETMAR in the U2OS, SM, SETF, MARF and SMF cell lines. The western blot was performed with anti-Hsmar1, anti-FLAG and anti-β-tubulin antibodies. The cell lines are described in Table 2.

**Table 2:** U2OS cell lines used in the *in vivo* DNA repair assay.

| Full name | Abbreviation | Expression level |
| --- | --- | --- |
| T-Rex-U2OS-TO | TO | Null |
| T-Rex-U2OS-SETMAR | SM2 | Low |
|  | SM3 | Very high |
| T-Rex-U2OS-TO-FLAG | TOF | Null |
| T-Rex-U2OS-SET-FLAG | SETF1 | Low |
|  | SETF2 | Medium |
| T-Rex-U2OS-SET-N210A-FLAG | SETF-N210A | Low |
| T-Rex-U2OS-MAR-FLAG | MARF | Medium |
| T-Rex-U2OS-SETMAR-FLAG | SMF2 | Medium |
|  | SMF3 | Medium |
| T-Rex-U2OS-SETMAR-N210A-FLAG | SMF-N210A | Medium |
| T-Rex-U2OS-SETMAR-R432A-FLAG | SMF-R432A | Medium |
| T-Rex-U2OS-SETMAR-D483A-FLAG | SMF-D483A | Medium |

The two control cell lines, TO and TOF, express only the endogenous SETMAR. We used four cell lines overexpressing wild type SETMAR at either low level, SM2, medium level, SMF2 and SMF3, or at very high level, SM3. The SET domain is overexpressed in SETF1 and SETF2 at low and medium level, respectively, whereas the MAR domain, which encodes the domesticated Hsmar1 transposase, is overexpressed at medium level. We also inserted three different mutations to abrogate specific functions of SETMAR. The N210A mutation, which is located in the key NHSC motif of the SET domain, abolishes the methyltransferase activity of SETMAR (7). To investigate the relative contribution of SETMAR binding to Hsmar1 transposon ends (inverted terminal repeat, ITR), we inserted the R432A mutation, which decreases the affinity of SETMAR to the Hsmar1 transposon ends (25, 27). To test the requirement of SETMAR’s trimming activity, we inserted the D483A mutation. The D483A mutant is catalytically defective because of the mutation of the first D of the DDD triad, which is necessary for the incorporation of one the Mg^2+^ ion (14, 25).

### The SET and MAR domains but not SETMAR promote DNA integration

SETMAR was previously shown to promote illegitimate integration in the genome. We used the different Flag-tagged SETMAR constructs to gain a better understanding of which SETMAR domains and activities are involved in DNA integration. For integration to happened, two conditions are required (Fig 3A). First, a plasmid need to be linearized by a DSB and it needs to be in the vicinity of a genomic DSB since integration is mediated by the NHEJ pathway (11). However, illegitimate integration is one of the three possible outcomes for a linearized plasmid because it can be either re-circularized or degraded (Fig 3A). The illegitimate plasmid integration rate was determined by transfecting a plasmid encoding a neomycin resistance marker before challenging the cells with G418 for two weeks. Cells in which the plasmid has been integrated into the genome could develop into foci. The foci were counted after staining with methylene blue.

**Fig 3.**
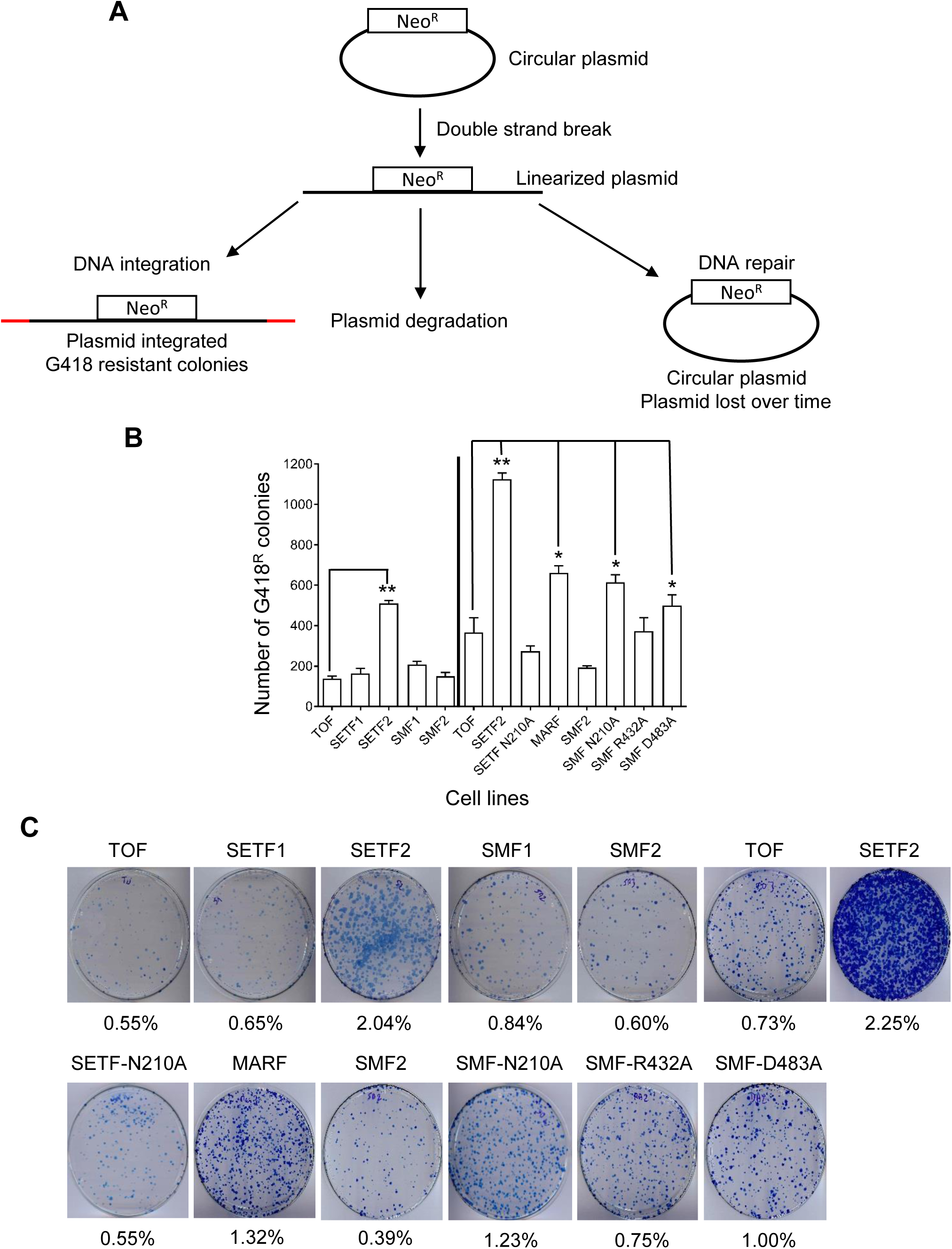
The SET and MAR domains increase the frequency of illegitimate DNA integration. **A**, Representation of the integration assay. Cells are transfected with a circular plasmid encoding a neomycin resistance gene. For integration to occur through the NHEJ pathway, the plasmid needs to be linearized by a DSB and a plasmid free end has to be in close vicinity of a genomic DSB. The linearized plasmid can also be repaired, which re-circularized the plasmid, or be degraded. Following G418 treatment for two weeks, surviving cells form foci which can be detected by methylene blue staining. **B**, Number of illegitimate integration events in the genome of a circular plasmid encoding a neomycin resistance gene. Average ± S.E.M. of 3 biological replicates. Statistical test: paired t-test, ^∗^ p-value < 0.05, ^∗∗^ p-value < 0.01. **C**, Representative pictures of integration plates. The integration rate for each cell line is indicated below each picture.

We first determined whether the topology of the plasmid influences its integration frequency. We therefore transfected a circular or linear version of the same plasmid in the control cell line TOF and observed a ~3-fold decrease in the integration of the linearized form compared to the circular plasmid (S1 Fig). We decided to use the circular plasmid for testing the different SETMAR constructs.

We performed two sets of plasmid integration experiments with the Flag-tagged cell lines (Fig 3B and C). A significant increase in illegitimate plasmid DNA integration was observed with a medium overexpression of the SET domain, the MAR domain and SETMAR N210A and R483A mutants. A low overexpression of the SET domain or medium overexpression of wild type SETMAR or SETMAR R432A mutant did not affect the plasmid integration rate. A representative selection of the integration plates of each cell line and their respective integration frequency are presented in Fig 3C. These results agree with published work showing that the efficiency of illegitimate recombination in most cell lines is less than 1% (11). We only observed an increase of the efficiency to 2% with a medium overexpression of the SET domain.

### The SET and MAR domains have an opposite effect on DNA repair

To gain a better understanding of SETMAR functions in the NHEJ pathway, we used a previously described *in vivo* DNA repair assay (28). This assay is based on two plasmids, one encoding the reporter gene (pEGFP) and the other serving as a transfection control (pRFP). The reporter plasmid encodes an *EGFP* gene interrupted by a 2.4 kb intron derived from the rat *Pem1* gene. An exon from the adenovirus (Ad2) has been integrated in the intron abolishing the GFP activity (Fig 4A). The Ad2 exon is flanked by HindIII and I-SceI restriction sites. Cleavage with HindIII or I-SceI yields compatible or incompatible ends, respectively (Fig 4B). These two types of ends require different steps for repair. Compatible ends can be directly ligated while incompatible ends need to be trimmed before the ligation step can occur. The repair of the linearized plasmid by the NHEJ pathway restores the GFP ORF making the cell green (Fig 4C). The repair events were detected by flow cytometry measuring at least 10,000 cells per assay. The repair efficiency was calculated as the ratio of green and red cells over the total number of red cells, thus normalizing the transfection efficiency between cell lines.

**Fig 4.**
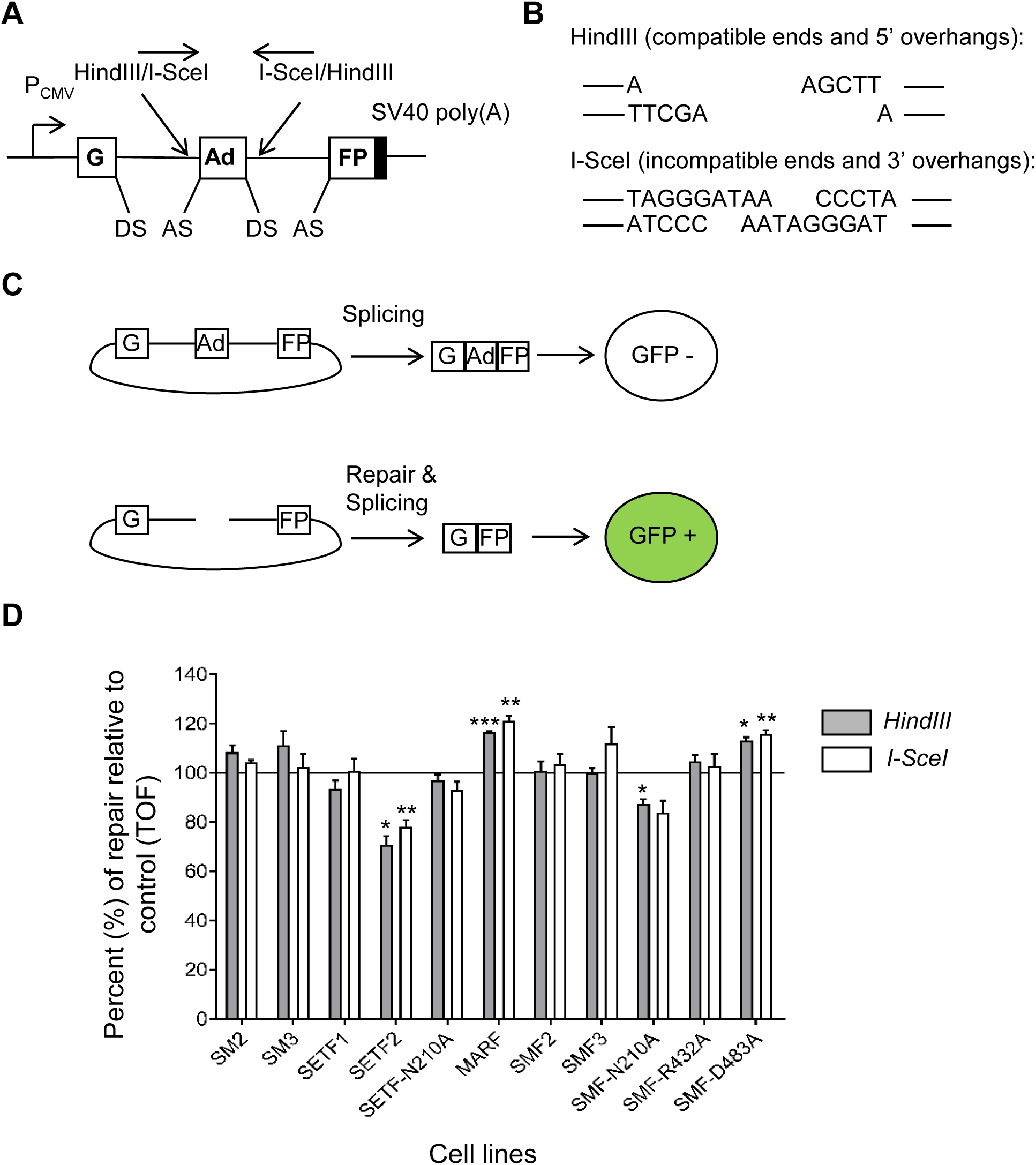
The SET and the MAR domains of SETMAR have an opposite effect on DNA repair by the NHEJ pathway. **A**, The reporter construct, pEGFP-Pem1-Ad2, is composed of a GFP cassette flanked by a PCMV promoter and a SV40 poly(A) sequence. The GFP coding sequence is interrupted by a 2.4 kb intron containing an adenovirus exon (Ad). The Ad exon is flanked by HindIII and I-SceI restriction sites. The donor (DS) and acceptor (AS) splicing sites are shown. **B**, HindIII and I-SceI restriction sites are respectively composed of a palindromic 6-bp and a non-palindromic 18-bp sequence. Digestion of the reporter construct by HindIII or I-SceI generates respectively compatible and incompatible ends. **C**, The presence of the Ad exon in the GFP ORF inactivates the GFP activity thus making the cell GFP negative. Removal of the Ad exon by HindIII or I-SceI followed by a successful intracellular repair will restore the GFP expression that can be quantified by flow cytometry. Adapted from (28). **D**, DNA repair efficiency of a linearized plasmid with compatible (HindIII) or incompatible (I-SceI) ends in the different cell lines relative to the control cell line (TO or TOF). Average ± S.E.M. of 3 biological replicates. Statistical test: paired t-test, ^∗^ p-value < 0.05, ^∗∗^ p-value < 0.01, ^∗∗∗^ p-value < 0.001.

For normalizing between the biological replicates, we calculated for each replicate the ratio of the repair efficiency of each cell line relative to their respective control cell line. The average ratio from three independent experiments is presented in Fig 4D. The SM2 and 3 cell lines were compared to the TO control cell line whereas the remaining cell lines were compared to TOF to take into account any possible effect of the Flag tag. Unsurprisingly, there was no significant difference in the repair efficiency of both types of ends between the two control cell lines (S2 Fig). In all cell lines, except SETF1 and 2, a ~10% decrease of the repair efficiency of incompatible ends compared to compatible ends was observed because of the necessary trimming of the DNA overhangs before the plasmid ends could be ligated (S2 Fig), in agreement with previously published results for this assay (28).

The overexpression of SETMAR, irrespective of the level of expression, does not promote the repair of either compatible or incompatible ends confirming the recent results obtained in the cell-extract assays (Fig 4D) (14). A slight but non-significant decrease in the repair of compatible ends is observed when the SET domain is expressed at a low level whereas a medium overexpression significantly decreases by ~20% the repair efficiency of both compatible and incompatible ends, suggesting a concentration-dependent effect. A low overexpression of the SETF N210A methyltransferase deficient mutant does not affect DNA repair of both type of ends. In contrast, medium overexpression of the MAR domain increases the repair efficiency of both types of ends by ~20% compared to the control cell line. The overexpression of SETMAR N210A reduces the repair efficiency of both type of ends by ~15% but only the compatible ends decrease is statistically significant, supporting a role for SETMAR methyltransferase activity for an efficient DNA repair (18). In contrast, SETMAR D483A mutant increases by ~15% the repair efficiency of both types of ends, which confirms that SETMAR nuclease activity is not required for DNA repair *in vivo* (14). No difference is observed with SETMAR R432A mutant for the repair of both types of ends confirming that the ITR binding activity is not involved in DNA repair.

## Discussion

It has been proposed that SETMAR could be involved in DNA repair through the NHEJ pathway. In this study, we investigated whether SETMAR, the SET or MAR domains and SETMAR methyltransferase, ITR binding and nuclease activities were involved in NHEJ and illegitimate DNA integration *in vivo*. We found that the SET and the MAR domains have an effect on DNA repair and integration but not wild type SETMAR. In addition, SETMAR proposed nuclease activity, which has been observed *in vitro*, does not seem to be functional *in vivo*. In contrast, SETMAR methyltransferase activity is required for an efficient DNA repair.

Previous publications have associated SETMAR expression to cell proliferation in different cell lines (22, 23). This association was not observed in the U2OS cell line (Fig 1B). Indeed, a modest overexpression of the SET domain or SETMAR only slightly reduces the growth rate after six to seven days. This could possibly be a non-specific effect, as protein overexpression per se is known to reduce growth rate (29). It is also possible that a strong reduction or increase in SETMAR expression is required to observe a change in cellular proliferation. Indeed, our cell lines have been selected for a modest overexpression of SETMAR rather than a strong one. Furthermore, we did not observe an enrichment in cell cycle related genes in the genes differentially expressed upon SETMAR modest overexpression (7).

The DNA repair and the DNA integration assays used in this study rely on the NHEJ pathway. The current model of plasmid integration is based on the group of proteins from classical NHEJ (11). This is supported by the idea that for the cell, DNA repair or DNA integration of a linear plasmid produces the same result, i.e. the removal of free ends, which could induce apoptosis or cell cycle arrest. The choice between the outcomes is dependent of the presence or not of a plasmid free end in the vicinity of a genomic DSB. The connection between DNA repair and DNA integration is also supported experimentally with cells depleted for a DNA repair protein which are unable to integrate DNA (30, 31).

Linearization of a plasmid is required before its integration in the genome (11). However, a linearized plasmid is more likely to be repaired than to be integrated. Indeed, the probability for the DNA repair complex to find the other plasmid end is higher than being near a genomic lesion due to physical continuity of the plasmid DNA, which ensures that the two ends can never be very far apart. Therefore, promotion of DNA repair could reduce the amount of linear plasmid in the cell by re-circularizing them and through it, decreasing the frequency of integration. The inefficiency of plasmid integration in mammalian cells comes from the combination of several limitations. The major ones are the low frequency of plasmid linearization, the re-circularization of the majority of linearized plasmids by the NHEJ pathway or their degradation, and the low probability of having a genomic and a plasmid end near each other. The proportion of linearized plasmid degraded by the cell is unknown but it is likely to be high. Indeed, the transfection of a linearized plasmid decreases by 3-fold the integration frequency compared to its circular form (S1 Fig). From the data obtained in our NHEJ assay we can estimate the frequency of re-circularization as at least 65% (S1 Fig). Since the integration frequency is below 1%, we can estimate that around 35% of the linear plasmid are degraded. A fraction of the transfected cells may also die from apoptosis if the free ends are not repaired in time. However, in the present experiment this is controlled by using a counted aliquot of the living cells for the G418 selection.

Previous works on the function of SETMAR in NHEJ claim specific roles for the SET and the MAR domains (18, 26). The SET domain dimethylates H3K36 of nucleosomes near DSBs. This epigenetic mark recruits and stabilizes the binding of Ku70 and NBS1 to the DNA ends (18). The MAR domain trims damaged and undamaged DNA overhangs before other NHEJ proteins ligate the ends (13). It has also been claimed that SETMAR activity is regulated by several interactions with other proteins involved in the NHEJ such as PRPF19 and DNA ligase IV (16, 17). Only a direct interaction between the SET domain and PRPF19 has been confirmed and is supposed to promote the recruitment of SETMAR to DSBs (16).

Overexpression of the wild type SETMAR did not affect DNA repair and integration in our *in vivo* assays. We found however that a medium overexpression of the SET domain, but not a low overexpression, decreases DNA repair efficiency and increases illegitimate DNA integration (Fig 3B and 4D). In our assays, both DNA repair and integration are supposed to be dependent on the NHEJ pathway, consistent with a role for the ancestral SET gene in this pathway. However, the mechanism by which the SET domain favours DNA integration over DNA repair is unclear. The decrease in recircularization with both compatible and incompatible ends found in the DNA repair assay could delay the re-circularization of plasmids, increasing the window of opportunity for a plasmid end to be in the vicinity of a genomic end and therefore promoting its genomic integration.

The overexpression of the MAR domain stimulates both DNA repair and integration (Fig 3B and 4D). These results seem to indicate that the proportion of linear plasmid degraded is reduced upon overexpression of the MAR domain. It has been previously proposed that the MAR domain of SETMAR could bind DNA free ends to trim the DNA overhangs (13, 14). However, if the MAR domain have a trimming activity, we would expect to observe a larger increase in the repair of incompatible ends versus compatible ends. In fact, the increase in the efficiency of repair is similar for both types of ends, which does not support a trimming activity (Fig 4D). Also, overexpression of SETMAR D483A mutant, which should abolish any remaining catalytic activity of the MAR domain, does not decrease the DNA repair efficiency (Fig 4D). In fact, we observe an increase in DNA integration and in DNA repair with both compatible and incompatible ends (Fig 3B and 4D). This seems to indicate that the MAR domain of SETMAR does not trim DNA overhangs *in vivo* but could however bind to DNA free ends to protect them from degradation, increasing the probability of integrating the linearized plasmid or re-circularizing the plasmid through the NHEJ pathway. The increase in DNA repair and integration could thus be mediated through the interactions of SETMAR with other NHEJ factors. It remains unknown whether these interactions are solely dependent on the SET domain or could also be mediated by the MAR domain. It is however important to remember that in our system the endogenous SETMAR is still expressed and the expression of the MAR domain could thus result in MAR dimers and in SETMAR-MAR dimers since SETMAR dimerization is mediated by the MAR domain. The SETMAR-MAR dimers could therefore be the only protein bringing the NHEJ factors to the DNA free ends if the interactions are dependent on the SET domain.

Unsurprisingly, the ITR binding activity of SETMAR is not required for DNA repair and integration (Fig 3B and 4D). In contrast, a medium overexpression of the methyltransferase defective mutant, SETMAR N210A, decreases DNA repair and increases DNA integration whereas overexpression of the wild type SETMAR does not affect DNA repair and integration (Fig 3B and 4D). The absence of effect of the N210A mutation in the SET domain construct is likely due to its low expression similar to SETF1, which does not affect DNA repair and integration. SETMAR N210A supports a role for the methyltransferase activity for an efficient DNA repair. It remains however unclear whether this is mediated by the deposition of H3K36me2 or by the methylation of another factors. Two studies, which also observe an increase in H3K36me2 at DSB sites, linked this increase to the removal of histone demethylases from the chromatin rather than active methylation (19, 20). In contrast, a recent study did not found any increase in H3K36me2 at a DSB site but found instead an increase in H3K36me3 (21). We recently found that SETMAR N210A mutant was decreasing the bulk level of H3K36me2 by western blot and also observed a decrease of H3K36me3 at some genomic positions, possibly because of a decreased H3K36me2 level which is required by SETD2 for adding the third methyl group (7). The decreased DNA repair activity with SETMAR N210A could therefore be due to this reduced H3K36me2/me3 level which would affect the repair efficiency by the NHEJ pathway. We must however stress that our analysis is based on a plasmid assay whereas previous observations were done on genomic DSBs.

An interesting question is why the SET and the MAR domains have an effect on DNA repair and integration but not the wild type SETMAR? A possibility is that the endogenous level of SETMAR in U2OS cells is already sufficient for an efficient DNA repair and therefore increasing wild type SETMAR level will not affect the NHEJ pathway. Another possibility is that SETMAR is a dimer in solution whereas almost all mammalian histone methyltransferase function as monomers (32, 33). The only known exception is vSET, a viral histone methyltransferase, which is active only as a dimer (34). The crystal structure of the SET domain is also a monomer strengthening the hypothesis that the pre-fusion SET gene was operating as a monomer (35). The MAR domain enforces the dimerization of SETMAR so even though the SET domain does not dimerize sensu stricto, the proximity between the two SET domains could however affect their methyltransferase activity or their interactions.

An interesting observation supporting this hypothesis is the presence of a SETMAR isoform encoding a defective histone methyltransferase monomer because of a splicing event which removes the majority of the SET and post-SET domains in the second exon. Interestingly, this SETMAR isoform is specific to the species where the SET gene is fused to the Hsmar1 transposase (Fig 5). The 5’ donor site is present in primates and other several mammals but the acceptor site is only found in anthropoid primates, except for the old-world monkeys where a single mutation in their common ancestor abolishes the acceptor site. The marmosets also lost their 5’ donor site but another less conserved site is present 20 nucleotides away. This isoform is expressed in most human tissues but at a lower level than the main isoform encoding the active methyltransferase monomer (36). This means that some SETMAR dimers should contain only one SET domain and could therefore function differently from SETMAR dimers with two SET domains.

**Fig 5.**
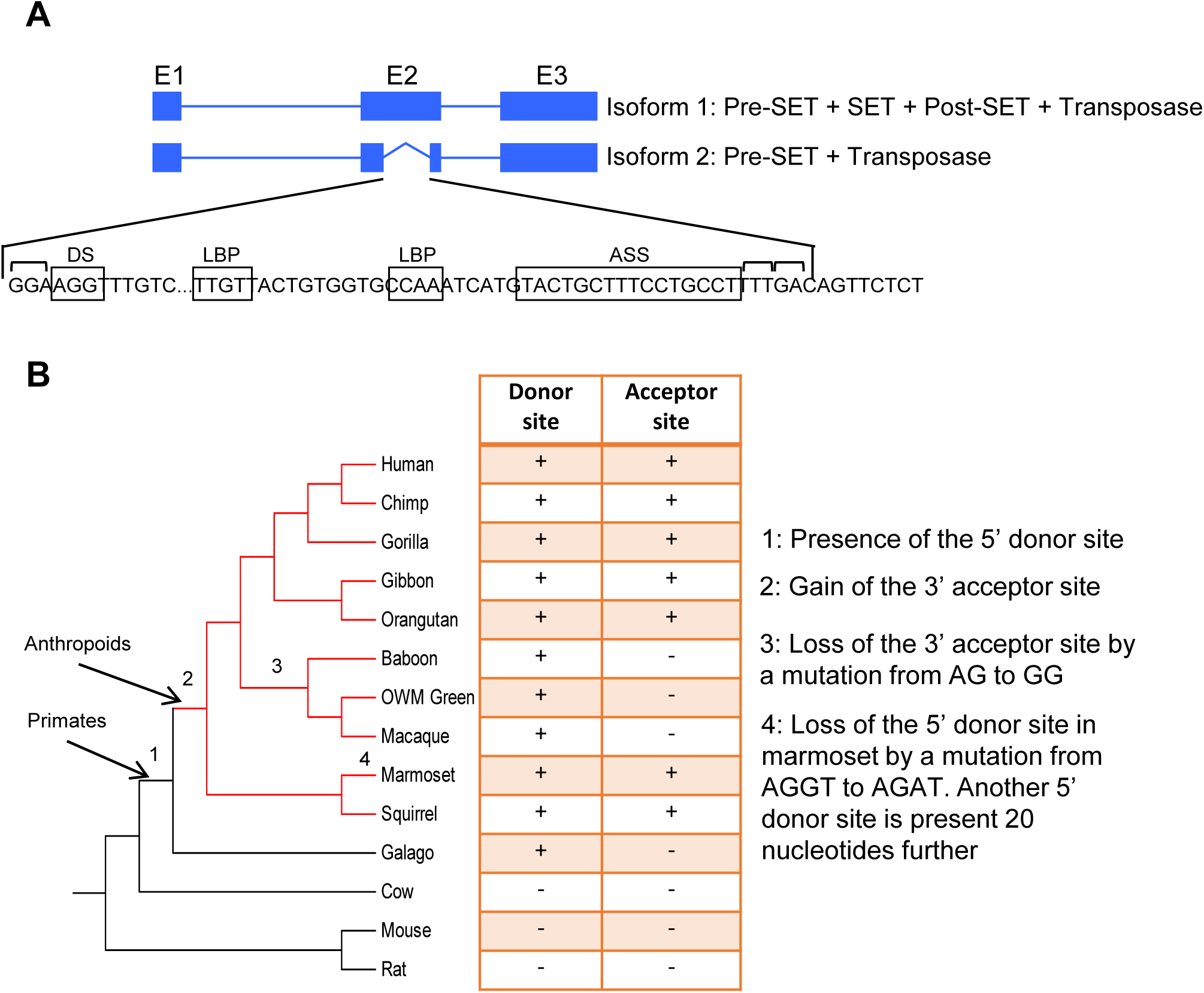
SETMAR second isoform is specific to the anthropoid primates. **A**, The human *SETMAR* gene encodes two major isoforms, the full-length protein (isoform 1) and a truncated protein (isoform 2). The second isoform is methyltransferase deficient because of the deletion of the majority of the SET and post-SET domains. Canonical donor site (DS), lariat branch points (LBP), and acceptor splicing site (ASS) are present in the second exon of SETMAR. The top brackets represents the exon codons. **B**, Phylogenetic tree of SETMAR second exon in several mammals. The 5’ donor site is found in all primates except for the marmoset (see 4) but should have appeared before the appearance of primates because of his presence in several non-primates mammals. The 3’ acceptor site is specific to anthropoid primates except for the old-world monkeys which lost it with a single point mutation.

## Author Contributions

Conceived and designed the experiments: MT. Performed the experiments: MT. Analyzed the data: MT. Contributed reagents/materials/analysis tools: MT RC. Wrote the paper: MT.

## Supporting Information

**S1 Fig. A circularized plasmid is more efficiently integrated than a linearized plasmid**. **A**, Comparison of the integration efficiency in the control cell line, TOF, of a circularized or linearized plasmid encoding a neomycin resistance gene. Average ± S.E.M of three biological replicates. Statistical test: paired t-test, ^∗∗^ p-value < 0.01. **B**, Representative pictures of integration plates. The integration rate of each cell line is indicated below each picture.

**S2 Fig. Representative FACS profiles of each cell lines**. FACS profiles of a pRFP plasmid used together with the pEGFP-Pem1-Ad2 reporter substrate are shown for each cell line. The profiles were generated using HindIII-(H3) and I-SceI-(SI) linearized plasmids. The proportion of repaired substrate is indicated in the lower right-hand corner of each profile. The percent is calculated from the number of cells that were doubly EGFP (horizontal) and RFP (vertical) positive versus the number of RFP positive.

